# Genetic mapping of some key plant architecture traits in *Brassica juncea* using a cross between two distinct lines – vegetable type Tumida and oleiferous Varuna

**DOI:** 10.1101/2022.07.11.499534

**Authors:** Shikha Mathur, Priyansha Singh, Satish Kumar Yadava, Vibha Gupta, Akshay K. Pradhan, Deepak Pental

## Abstract

*Brassica juncea* (AABB, 2n=36), commonly called mustard is an allopolyploid crop of recent origin but has considerable morphological and underlying genetic variation. An F_1_-derived doubled haploid (F_1_DH) population developed from a cross between a Indian oleiferous line, Varuna, and a Chinese stem type vegetable mustard, Tumida showed significant variability for some key plant architectural traits, including four stem strength-related traits, stem diameter, plant height, branch initiation height, number of primary branches (*Pbr*), and days to flowering (*Df*). Multi-environment QTL analysis identified twenty environmentally stable QTL for the nine plant architectural traits. Both Tumida and Varuna contain some positive QTL that can be used to breed superior ideotypes in mustard. A QTL cluster on LG A10 contained environmentally stable QTL for seven architectural traits. This region also contained overlapping environmentally stable major QTL (phenotypic variance ≥ 10%) for *Df* and *Pbr*, with Tumida contributing the trait enhancing alleles for both the traits. Since early flowering is critical for the cultivation of mustard in the Indian subcontinent, this QTL cannot be used for the improvement of *Pbr* in the Indian gene pool lines. Conditional QTL analysis for *Pbr* identified the QTL for improvement of *Pbr* without negative effects on *Df*. The environmentally stable QTL were projected on the genome assemblies of Tumida and Varuna for the identification of candidate genes. These findings provide insights into the genetics of plant architectural traits in two diverse gene pools of *B. juncea* and provide opportunities for improvement of the plant architecture through marker-assisted introgressions.

**Key Message:** Genetic mapping of some key plant architectural traits in a vegetable type and an oleiferous *B. juncea* cross revealed environmentally stable QTL and candidate genes for breeding more productive ideotypes.

## INTRODUCTION

*Brassica juncea* (L.) Czern & Coss (AABB, 2n=36) is a major oilseed crop of the Indian subcontinent cultivated in more than six million hectares of land in India alone (Jat et al. 2019). The crop is well-adapted for cultivation in dryland areas but requires genetic improvement for higher yield and resistance to pests. The yield increase in a crop, either by varietal or hybrid development or through yield protection, depends upon the genetic variability available within the crop species. We earlier showed the presence of two distinct gene pools amongst the oleiferous types of *B. juncea* – the Indian gene pool and the east European gene pool; this identification was based on the differences in morphological and reproductive traits (Pradhan et al. 1993), and on a molecular marker-based diversity analysis (Srivastava et al. 2001). The Indian and the east European germplasm lines have been extensively used for genetic mapping of many traits of high agronomic value like disease resistance (Panjabi-Massand et al. 2010; Arora et al. 2019), oil content (Rout et al. 2018), yellow seed coat color (Padmaja et al. 2014), seed size (Dhaka et al. 2017) and several other yield influencing traits (Ramchiary et al. 2007; Yadava et al. 2012). The hybrids between the Indian and the east European gene pool lines of *B. juncea* were found to be heterotic for yield (Pradhan et al. 1993).

Although, *B. juncea* is a recent allopolyploid – extensive diversity has been reported within its primary gene pool. A recent genome sequencing of 480 lines of *B. juncea* has revealed six distinct genetic groups within the primary gene pool of the species (Kang et al. 2021). Two of these genetic groups, G2 and G6 have been earlier described as the east European gene pool (G2) and the Indian gene pool (G6) (Srivastava et al. 2001). Four additional distinct genetic groups (G1, G3, G4, and G5) have also been identified amongst the *B. juncea* lines under cultivation predominantly in east Asia. We will refer to the genetic groups as gene pools because these groups have undergone differential selection pressure due to their geographical location and end-use; however, they are a part of the primary gene pool of *B. juncea*. A *B. juncea* vegetable type stem mustard line Tumida (G5) was the first *B. juncea* genotype to be sequenced (Yang et al. 2016). We recently reported a highly contiguous genome assembly of an oleiferous type of *B. juncea* variety Varuna belonging to the Indian gene pool (G6) (Paritosh et al. 2021). Tumida types are distinct from lines of the Indian oleiferous gene pool like Varuna for plant architecture and yield influencing reproductive traits. The most distinct feature of the Indian gene pool lines (G6) is that they do not have any photoperiod sensitivity or vernalization requirements for flowering unlike the lines of other gene pools (G1-G5). The Indian gene pool lines are grown during a short winter period in the northern plains of India whereas, the other gene pool lines are grown during the summer season under long day-length conditions and therefore, are ill-adapted to the growing conditions in the Indian subcontinent.

Modification of plant architecture has been shown to contribute to the improvement of yield and/or stress resistance in several crops (Guo et al. 2020). As an example, the mustard hybrids developed by our group have a significantly large number of primary and secondary branches and high pod density (Sodhi et al. 2006; Aakanksha et al. 2021) but are tall and therefore, susceptible to breaking and lodging under high winds.

An F_1_DH (F_1_-derived doubled haploid) population developed from a cross between Tumida and Varuna (hereinafter referred to as the “TUV” population), showed striking variability for several plant architectural traits. We report here the mapping of some crucial plant architectural traits – stem strength and diameter, plant height, branch initiation height, number of primary branches, and days to flowering which would be useful in developing lines with improved lodging resistance, reduced plant height, branch initiation height, and days to flowering and higher number of primary branches. The results clearly show that exotic germplasm, like Tumida, seemingly ill-adapted to the Indian growing conditions, can contribute some significant QTL to improve lines of the Indian and other gene pools of *B. juncea* for some important architectural traits.

## MATERIALS AND METHODS

### Plant materials, field experiments, and phenotypic evaluation

*B. juncea* line Tumida was crossed with line Varuna, and F_1_ plants were used for developing DH lines by microspore culture following an earlier described protocol (Mukhopadhyay et al. 2007). Both the lines – Tumida and Varuna had been maintained by strict selfing. For phenotyping, the parents, F_1_, and the 169 F_1_DH lines were grown in four independent trials (T1, T2, T3, and T4) at the University of Delhi research farm station in Bawana, Delhi, India during the crop growing seasons of 2018-19 (T1), 2019-20 (T2) and 2020-21 (T3 and T4 – staggered sowing). The TUV population was sown in a randomized complete block design with three replications in each trial. Each line was planted in a single 2 m row with a 45 cm distance between two adjacent rows; the sowing density was ~20 plants per row. More details on the field trials are provided in Table S1.

Phenotyping was carried out for several traits which included, stem strength-related traits, stem diameter (*Dia*), branch initiation height (*Bih*), plant height (*Plht*), number of primary branches (*Pbr*), and days to flowering (*Df*). For *Bih, Plht*, and *Pbr*, data was taken from three competitive plants from each replication and the mean of the observations was used as the trait value. *Bih* was measured as the length from the base of the plant to the point from where the first primary branch arises. *Df* was recorded when about 50% of the plants in a row had at least one flower open. For measurements of stem strength-related traits and stem diameter (*Dia*), the stem tissues were sampled at the stage when seed filling ended but pods were still green, and the stem bore maximum weight (hereinafter referred to as the “mature green stage”). A total of ten competitive plants from each replication were sampled and the mean of the observations was used as the trait value. The measurements for these traits were taken at the mid-point of the last internode immediately below the inflorescence (hereinafter referred to as the “test-point”). *Dia* was measured using the digital vernier calipers (Model No. 1108-150, INSIZE Co. Ltd., India). In the case of the stem strength-related traits – bend force (*Bf*) was measured using a three-point bending test, stab force (*Sf*) was measured using a pinhead to penetrate the stem epidermal layers and vasculature, and press force (*Pf*) was measured using a flat-head to press the stem at the test-point. All stem strength-related traits (*Bf, Sf*, and *Pf*) were measured with a YYD-1 instrument (Zhejiang Top Cloud-Agri Technology Co. Ltd., Hangzhou, China). Stem breaking strength (*Bs*) was calculated as described by Wei et al. (2017).

The Best Linear Unbiased Prediction (BLUP) for each TUV line across the three environments was computed for each of the nine traits using the R package ‘metan’ (Olivoto and Lúcio 2020) with a mixed linear model that accounts for the effects of the environment, replication, genotype, and genotype by environment. The average trait values obtained in each trial – single environment (SE) values and BLUPs were subjected to correlation statistical analyses using Plabstat (Utz 2001). The broad-sense heritability (H_B_) for each trait was computed using the phenotyping module of iMAS (Integrated Marker Assisted Selection) version 2.1 (Sirisha et al. 2005) using the average trait values obtained in each field trial.

### Genetic mapping, QTL analysis, and identification of the candidate genes

The 169 TUV lines were used for the construction of a genetic map using GBS (genotyping by sequencing) based SNP markers that have been described earlier (Paritosh et al. 2021). The genetic map was developed using the ‘mstmap’ function of the ASMap package in R (Taylor and Butler 2017) using the parameters – distance function Kosambi, cut-off *p* value 1e-15, and a missing threshold of 0.3.

QTL mapping was carried out with the SE trait values and BLUPs using the composite interval mapping (CIM) module of Windows QTL Cartographer 2.5 (Wang 2007). For CIM, the standard model (Model 6) was used with forward regression, a window size of 10 cM, and five background control markers. The genome was scanned for putative QTL with main effects at a walking speed of 1 cM. The experiment-wise error rate was determined by performing 1000 permutations to obtain the empirical thresholds (Churchill and Doerge 1994). A LOD threshold of 3.0 was used for identifying significant QTL. A QTL with LOD values ranging between 2.5 and 3 was considered a suggestive QTL. QTL detected with phenotypic variance (R^2^) ≥ 10% was considered a major QTL.

The QTL were named using an abbreviation of the trait beginning with a capital letter followed by the name of the linkage group (LG) and the serial number of chronological QTL for the trait detected on the LG independently in each analysis. The QTL detected using SE trait values were superscripted with the abbreviation for the trial (T1, T2, T3, and T4) in which these were detected. The QTL detected using BLUPs were superscripted with the letter ‘B’. The QTL identified for a trait in different analyses with overlapping confidence intervals were assumed to be the same QTL. A QTL that could be detected in at least two single environments (SEs) and also using BLUPs was regarded as a “Stable QTL” as described earlier (Xu et al. 2022). The Stable QTL was prefixed with ‘S-’ and the confidence interval of the QTL detected using BLUPs was used as the confidence interval for the Stable QTL.

Epistatic QTL were detected using QTL Network 2.0 (Yang et al. 2007). The program simultaneously detects the additive QTL, QTL × environment (QE), epistasis, and epistasis × environment interactions using multi-environment trait data. The additive QTL detected using QTL Network 2.0 were suffixed with ‘.QN’. The analyses were carried out using mixed-model-based composite interval mapping (MCIM) with 1 cM walk speed and a testing window of 10 cM. Thresholds for the presence of QTL were generated by performing 1000 permutations. The epistatic effects among loci with or without individual additive main effects were detected by performing 2D genome scans. The digenic interactions (additive × additive) studied included interactions between two main-effect QTL (Type I interaction), between a main effect QTL and a QTL without any significant main effect (Type II interaction), and between two QTL without any significant main effect (Type III interaction) (Li et al. 2001).

Conditional QTL mapping was carried out using the software QGA Station 2.0 (http://ibi.zju.edu.cn/software/qga/v2.0/index.htm; Zhu 1995). QTL mapping for conditional values was performed using Windows QTL Cartographer 2.5 as described above and the identified QTL were defined as conditional QTL superscripted with the letter ‘C’.

The GBS markers flanking the QTL regions were mapped on the genome assemblies of *B. juncea* lines Varuna (Paritosh et al. 2021) and Tumida (Yang et al. 2016) using blastn, *e* value 1e-25, >95% identity, and 100% coverage. The sequence tags for the GBS markers used in the study have been provided earlier (Paritosh et al. 2021). The genes between the flanking markers and the annotations of Tumida-specific genes were based on the revised assembly of the Tumida genome (V2.0) available in the *Brassica* database (http://brassicadb.cn/#/; Chen et al. 2022). The GO annotations and functional classification of Varuna genes were obtained using the Blast2GO program (https://www.biobam.com/omicsbox; Conesa et al. 2005).

### Stem anatomy

The anatomy of the stem tissues was studied at the mature green stage. Transverse free-hand sections obtained at the test point from three different individuals were analyzed for each TUV line. A Leica TCS SP8 AOBS confocal laser scanning microscope (Leica Microsystems Mannheim, Germany) was used to examine the sections for lignin autofluorescence and collection of Differential Interference Contrast (DIC) images. The sections were mounted in 50% (v/v) glycerol for visualization of total lignin autofluorescence (excitation at 405 nm and emission detected at 440-510 nm). The laser intensity, pinhole, and photomultiplier gain settings were kept constant between samples to obtain meaningful comparisons.

## RESULTS

### Construction of the TUV linkage map

A linkage map of TUV F_1_DH lines using 9041 polymorphic genetic markers has been described earlier (Paritosh et al. 2021). In the present study, a subset of 2028 polymorphic GBS-based SNP markers was identified from the earlier TUV genetic map by removing the excess markers mapping to the same positions. These 2028 markers were used for developing a new linkage map using 169 individuals of the TUV population. The genetic map is 3512.8 cM in length with an average of 113 markers per linkage group at an average spacing of 1.73 cM between two consecutive markers. Information on the distribution, density, and positions of the markers on the TUV linkage map is provided in Tables S2 and S3.

### Phenotypic evaluation of the parents, F_1_, and the TUV population

The mean trait values of the parental lines, F_1_, and the range and mean trait values of the DH lines for the nine plant architectural traits namely, press force (*Pf*), bend force (*Bf*), stab force (*Sf*), breaking strength (*Bs*), stem diameter (*Dia*), plant height (*Plht*), branch initiation height (*Bih*), number of primary branches (*Pbr*), and days to flowering (*Df*) are listed in Table 1 and Table S4. The BLUPs of the TUV DH lines are provided in Table S5. Varuna showed higher trait values for the stem strength-related traits (*Pf, Sf, Bf*, and *Bs), Dia, Bih*, and *Plht* whereas Tumida showed higher trait values for *Pbr* and *Df* (Fig. 1 and Table 1). The low trait values for *Plht, Bih*, and *Df* are desirable from the breeding perspective; therefore, Tumida is the better parent for *Plht* and *Bih*, and Varuna is the better parent for *Df*. For most of the traits, the F_1_ showed intermediate values with the exceptions of *Dia*, *Plht*, and *Bih* (Fig. S1 and Table 1). The TUV mapping population showed transgressive segregation, suggesting that both Varuna and Tumida contained positive alleles for all the nine architectural traits (Fig. S1 and Table 1). However, very few transgressive segregants were observed for *Df* with a flowering time less than that of Varuna (Table 1 and Table S5). Extensive transgressive segregation was observed for the *Bih* trait with values ranging from 17.05-143.09 cm (Fig. S1 and Table 1).

**Fig. 1.**
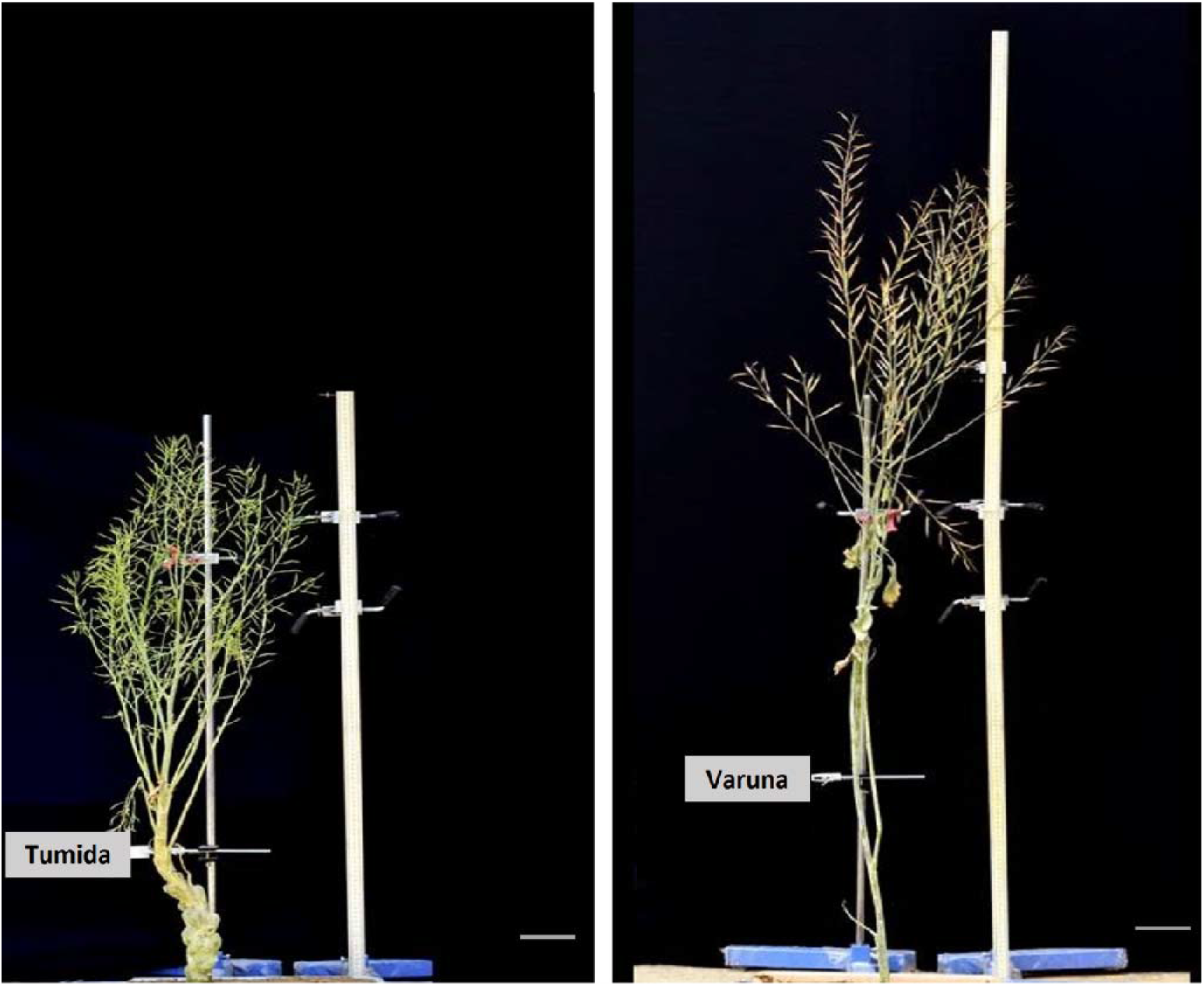
Morphotypes of Tumida and Varuna plants showing differences in their architecture at maturity

**Table 1.** Average trait values of Tumida, Varuna, and TUV-F_1_ and trait statistics on adjusted mean values (BLUP values) of the TUV population for the nine plant architectural traits

| Trait | Mean | | | F<sub>1</sub>DH population range | F<sub>1</sub>DH population mean ± S.D. |
| --- | --- | --- | --- | --- | --- |
|  | Tumida | Varuna | TUV-F<sub>1</sub> |  |  |
| *Bf* | 10.7 | 16.64 | 13.78 | 6.5-27.35 | 14.82 ± 3.5 |
| *Sf* | 9.23 | 20.64 | 17.84 | 13.29-27.05 | 16.95 ± 1.95 |
| *Pf* | 98.93 | 113.57 | 110.75 | 80.57-127.72 | 103.82 ± 9.61 |
| *Bs* | 0.54 | 1.43 | 1.39 | 0.8-2.09 | 1.27 ± 0.16 |
| *Dia* | 3.84 | 3.95 | 3.55 | 2.88-4.88 | 3.76 ± 0.37 |
| *Plht* | 134.32 | 180.36 | 208.4 | 154.25-256.11 | 210.55 ± 23.11 |
| *Bih* | 38.77 | 51.42 | 89.98 | 17.05-143.09 | 76.71 ± 30.16 |
| *Pbr* | 10.26 | 5.03 | 8.82 | 2.99-13.43 | 7.04 ± 2.19 |
| *Df* | 131.12 | 42.8 | 77.51 | 42.77-123.13 | 79.81 ± 21.49 |

*Bf*, *Sf*, *Pf*, *Bs*, *Dia*, *Plht*, *Bih*, *Pbr*, and *Df* represent bend force, stab force, press force, breaking strength, stem diameter, plant height, branch initiation height, number of primary branches, and days to flower, respectively.

The broad sense heritability (H_B_) of the nine plant architectural traits ranged from 33 to 98% in the three environments (Table S4). The stem strength-related traits were found to be moderately heritable with H_B_ of *Pf, Sf, Bf*, and *Bs* ranging from 33-36%, 45-60%, 48-62%, and 42-62%, respectively (Table S4). *Dia* and *Plht* also showed moderate heritability of 48-62% and 53-89%, respectively. *Bih* (H_B_ 79-88%), *Pbr* (H_B_ 81-90%), and *Df* (H_B_ 94-98%) were found to be highly heritable traits.

We also undertook an anatomical analysis of the stems of Tumida, Varuna, and some TUV population lines (TUV-547, TUV-780, TUV-95, and TUV-25) showing transgressive segregation for most of the stem strength-related traits (Fig. 2 and Fig. S2). The transverse sections of stems of parents at the test-point revealed more lignified interfascicular sclerenchyma tissue in Varuna as compared to Tumida (Fig. 2 and Fig. S2), suggesting differential cambial activity in Tumida and Varuna. Similar differences were also observed in the TUV lines that showed contrasting values for the stem strength-related traits (Fig. S2).

**Fig. 2.**
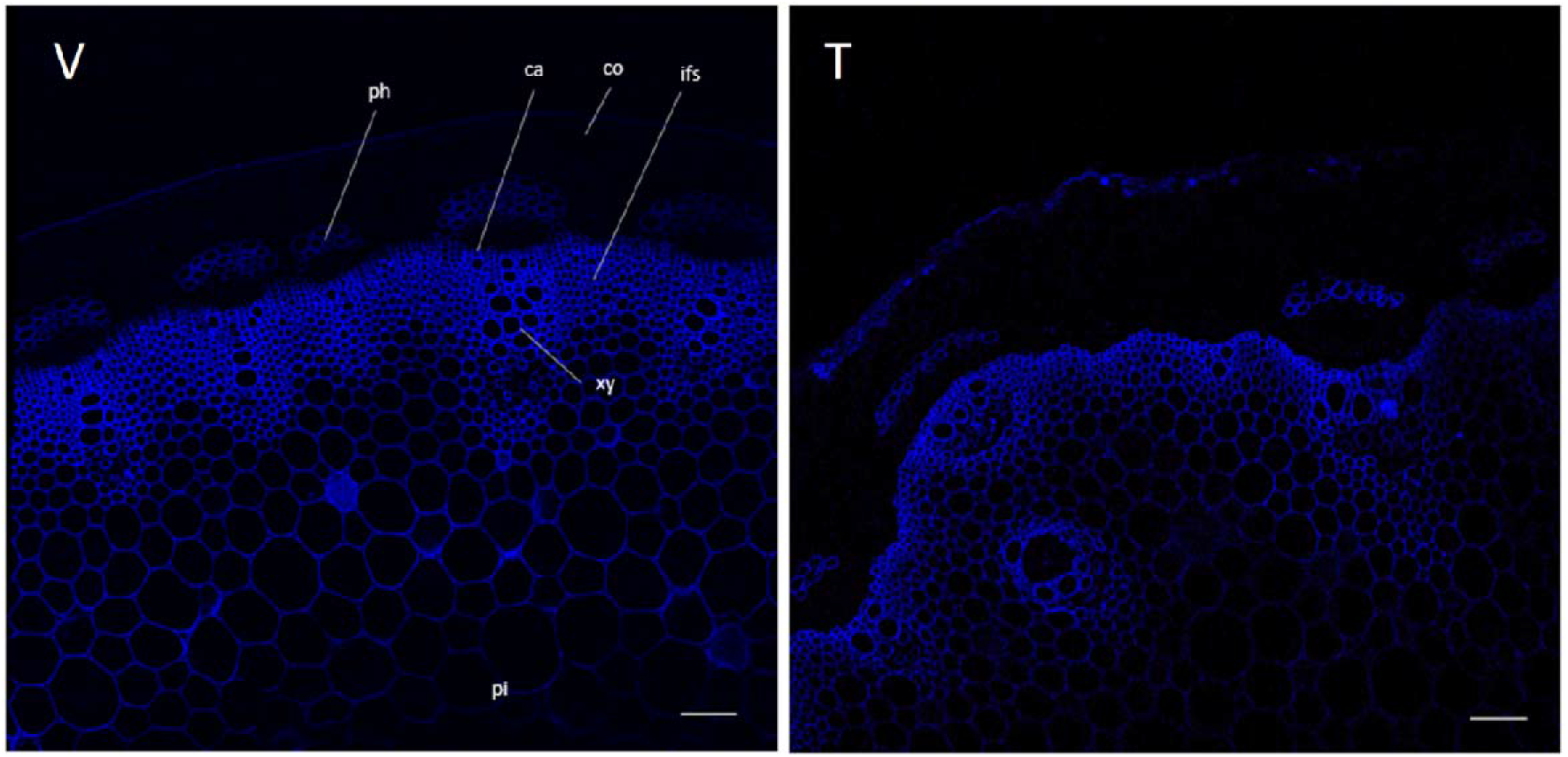
Stem sections of Varuna (V) and Tumida (T) showing blue UV autofluorescence from lignin. Scale bars represent 100 μm. Abbreviations: co, cortex; xy, xylem; ph, phloem; ca, cambium; ifs, interfascicular sclerenchyma; pi, pith

### Correlation analysis of the plant architectural traits

The Pearson correlation coefficients among the nine quantitative traits were estimated using BLUPs (Table 2). The four stem strength-related traits (*Pf, Sf, Bf*, and *Bs*) showed a significantly (*p* ≤ 0.01) high positive correlation with one another. *Dia* showed a significant (*p* ≤ 0.01) positive correlation with *Bf* and *Pf* whereas, it was not highly correlated with *Bs* and *Sf. Bih* and *Plht* were positively correlated with one another. *Df* showed a significant (*p ≤* 0.01) negative correlation with *Pf, Sf, Bf*, and *Dia* and a positive correlation with *Plht* and *Bih. Pbr* showed a significant (*p* ≤ 0.01) positive correlation with *Df* (r = 0.737) and *Plht* (r = 0.668) which might negatively affect the simultaneous improvement of these traits.

**Table 2.** The Pearson correlation coefficients amongst the adjusted mean (BLUP) values of the nine plant architectural traits

|  | *Pf* | *Sf* | *Bf* | *Bs* | *Dia* | *Plht* | *Bih* | *Pbr* |
| --- | --- | --- | --- | --- | --- | --- | --- | --- |
| *Sf* | 0.607** |  |  |  |  |  |  |  |
| *Bf* | 0.824** | 0.713** |  |  |  |  |  |  |
| *Bs* | 0.568** | 0.826** | 0.674** |  |  |  |  |  |
| *Dia* | 0.728** | 0.386** | 0.847** | 0.222** |  |  |  |  |
| *Plht* | -0.058 | 0.082 | -0.075 | 0.215** | -0.219** |  |  |  |
| *Bih* | -0.269** | -0.102 | -0.308** | 0.039 | -0.437** | 0.832** |  |  |
| *Pbr* | -0.356** | -0.075 | -0.351** | 0.095 | -0.548** | 0.668** | 0.779** |  |
| *Df* | -0.497** | -0.403** | -0.537** | -0.226** | -0.591** | 0.592** | 0.837** | 0.737** |

\*significant at 5%
\*\*significant at 1%
*Bf*, *Sf*, *Pf*, *Bs*, *Dia*, *Plht*, *Bih*, *Pbr*, and *Df* represent bend force, stab force, press force, breaking strength, stem diameter, plant height, branch initiation height, number of primary branches, and days to flower, respectively.

### QTL Mapping

The QTL for each trait were detected using Windows QTL Cartographer 2.5 using the mean trait values obtained in each trial (SE trait values) (Table S6) and using the BLUPs (Table S7). A QTL that could be detected in at least two single environments (SEs) and also detected using BLUPs, was designated a Stable QTL. QTL mapping was also carried out using QTL Network 2.0, which allowed the detection of QTL × environment (QE), epistasis, and epistasis × environment interactions in addition to the detection of additive QTL (Tables S8 and S9). An additive QTL detected for a trait by both Windows QTL Cartographer 2.5 and QTL Network 2.0 with overlapping confidence intervals was considered to be the same QTL (Table S8). For every trait, we first describe the QTL mapped using Windows QTL Cartographer 2.5, followed by an analysis with QTL Network 2.0.

#### Stem strength-related traits (*Pf, Sf, Bf*, and *Bs*)

A total of 13, 18, 15, and 12 QTL were detected using SE trait values for *Pf, Sf, Bf*, and *Bs*, respectively in the three trials (Table S6). Using BLUPs, four QTL each for *Pf, Bf*, and *Bs* and three QTL for *Sf* were detected (Table S7). Several QTL for *Pf, Sf, Bf*, and *Bs* mapped to overlapping intervals, particularly, on linkage groups (LGs) A10 and B07. This observation parallels the significantly high positive correlations predicted between these traits. A total of seven Stable QTL were identified for stem strength-related traits (*Pf, Sf*, and *Bf*) on LGs A07, A09, A10, B06, and B07 (Fig. 4, Table 3, and Table S10). Varuna contributed beneficial alleles for the major Stable QTL for *Pf* (*S-Pf-A10-1*) and *Bf* (*S-Bf-A10-1*) in overlapping confidence intervals on LG A10. These QTL explain 23% and 19.5% of the phenotypic variances, respectively. A Stable QTL *S-Pf-B07-1*, explaining 7.1% of the phenotypic variance was contributed by Tumida. We detected three Stable *Sf* QTL – *S-Sf-A09-1, S-Sf-A10-1*, and *S-Sf-B06-1* with Varuna contributing positive alleles for the trait (Fig. 4, Table 3, and Table S10). No Stable QTL was detected for *Bs* using the adopted criteria.

**Fig. 3.**
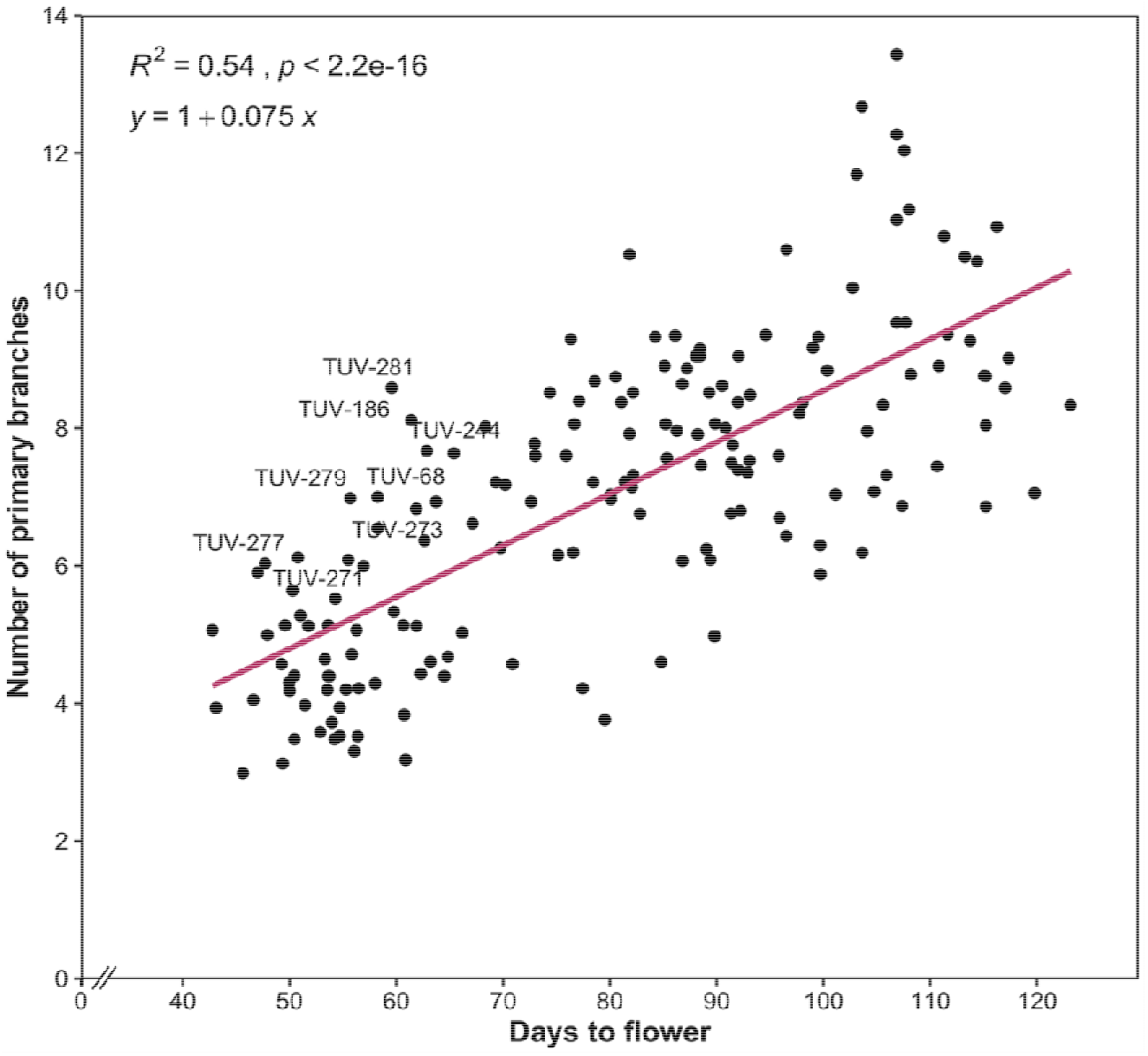
Regression of the number of primary branches (*Pbr*) trait on days to flowering (*Df*) in the TUV population

**Fig. 4.**
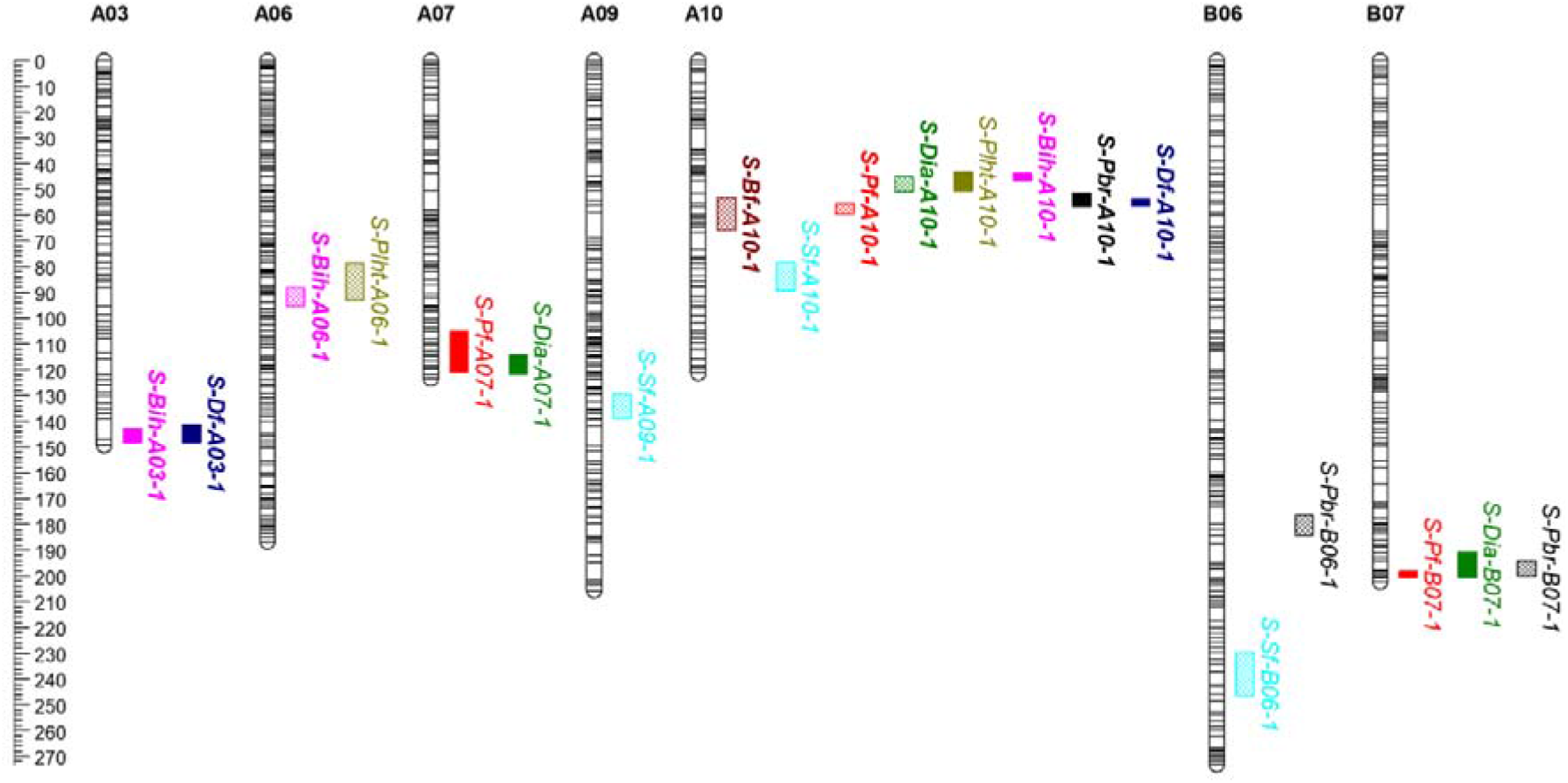
Genetic locations of Stable QTL regions associated with press force (*Pf*), stab force (*Sf*), bend force (*Bf*), breaking strength (*Bs*), stem diameter (*Dia*), plant height (*Plht*), branch initiation height (*Bih*), number of primary branches (*Pbr*) and days to flower (*Df*). Uniform cM scale is shown on the left. Solid and checked bars represent the QTL from Tumida and Varuna, respectively. Major QTL (R^2^ ≥ 10%) are highlighted in bold

**Table 3.**
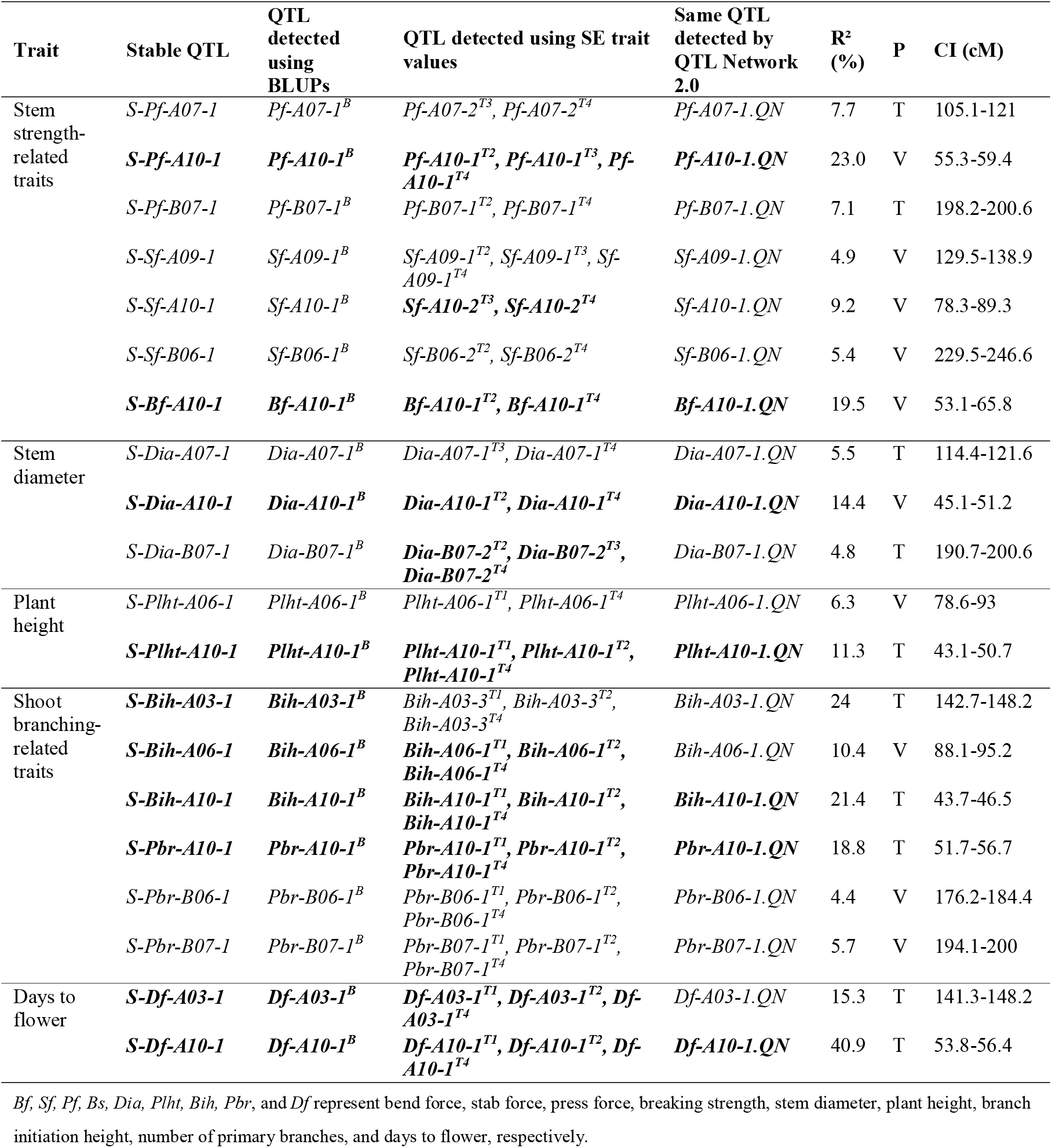
The Stable QTL detected for plant architectural traits in the TUV population. Major QTL (R^2^ ≥ 10%) have been highlighted in bold. BLUP, Best Linear Unbiased Prediction; SE, single environment; T1, trial 1; T2, trial 2; T3, trial 3; T4, trial 4; R^2^, phenotypic variance explained by the QTL; P, parent contributing the trait enhancing allele; T, Tumida; V, Varuna; CI, confidence interval

| Trait | Stable QTL | QTL detected using BLUPs | QTL detected using SE trait values | Same QTL detected by QTL Network 2.0 | $R^2$ (%) | P | CI (cM) |
| --- | --- | --- | --- | --- | --- | --- | --- |
| Stem strength-related traits | *S-Pf-A07-1* | *Pf-A07-1<sup>B</sup>* | *Pf-A07-2<sup>T3</sup>, Pf-A07-2<sup>T4</sup>* | *Pf-A07-1.QN* | 7.7 | T | 105.1-121 |
|  | ***S-Pf-A10-1*** | ***Pf-A10-1<sup>B</sup>*** | ***Pf-A10-1<sup>T2</sup>, Pf-A10-1<sup>T3</sup>, Pf-A10-1<sup>T4</sup>*** | ***Pf-A10-1.QN*** | 23.0 | V | 55.3-59.4 |
|  | *S-Pf-B07-1* | *Pf-B07-1<sup>B</sup>* | *Pf-B07-1<sup>T2</sup>, Pf-B07-1<sup>T4</sup>* | *Pf-B07-1.QN* | 7.1 | T | 198.2-200.6 |
|  | *S-Sf-A09-1* | *Sf-A09-1<sup>B</sup>* | *Sf-A09-1<sup>T2</sup>, Sf-A09-1<sup>T3</sup>, Sf-A09-1<sup>T4</sup>* | *Sf-A09-1.QN* | 4.9 | V | 129.5-138.9 |
|  | *S-Sf-A10-1* | *Sf-A10-1<sup>B</sup>* | ***Sf-A10-2<sup>T3</sup>, Sf-A10-2<sup>T4</sup>*** | *Sf-A10-1.QN* | 9.2 | V | 78.3-89.3 |
|  | *S-Sf-B06-1* | *Sf-B06-1<sup>B</sup>* | *Sf-B06-2<sup>T2</sup>, Sf-B06-2<sup>T4</sup>* | *Sf-B06-1.QN* | 5.4 | V | 229.5-246.6 |
|  | ***S-Bf-A10-1*** | ***Bf-A10-1<sup>B</sup>*** | ***Bf-A10-1<sup>T2</sup>, Bf-A10-1<sup>T4</sup>*** | ***Bf-A10-1.QN*** | 19.5 | V | 53.1-65.8 |
| Stem diameter | *S-Dia-A07-1* | *Dia-A07-1<sup>B</sup>* | *Dia-A07-1<sup>T3</sup>, Dia-A07-1<sup>T4</sup>* | *Dia-A07-1.QN* | 5.5 | T | 114.4-121.6 |
|  | ***S-Dia-A10-1*** | ***Dia-A10-1<sup>B</sup>*** | ***Dia-A10-1<sup>T2</sup>, Dia-A10-1<sup>T4</sup>*** | ***Dia-A10-1.QN*** | 14.4 | V | 45.1-51.2 |
|  | *S-Dia-B07-1* | *Dia-B07-1<sup>B</sup>* | ***Dia-B07-2<sup>T2</sup>, Dia-B07-2<sup>T3</sup>, Dia-B07-2<sup>T4</sup>*** | *Dia-B07-1.QN* | 4.8 | T | 190.7-200.6 |
| Plant height | *S-Plht-A06-1* | *Plht-A06-1<sup>B</sup>* | *Plht-A06-1<sup>T1</sup>, Plht-A06-1<sup>T4</sup>* | *Plht-A06-1.QN* | 6.3 | V | 78.6-93 |
|  | ***S-Plht-A10-1*** | ***Plht-A10-1<sup>B</sup>*** | ***Plht-A10-1<sup>T1</sup>, Plht-A10-1<sup>T2</sup>, Plht-A10-1<sup>T4</sup>*** | ***Plht-A10-1.QN*** | 11.3 | T | 43.1-50.7 |
| Shoot branching-related traits | ***S-Bih-A03-1*** | ***Bih-A03-1<sup>B</sup>*** | *Bih-A03-3<sup>T1</sup>, Bih-A03-3<sup>T2</sup>, Bih-A03-3<sup>T4</sup>* | *Bih-A03-1.QN* | 24 | T | 142.7-148.2 |
|  | ***S-Bih-A06-1*** | ***Bih-A06-1<sup>B</sup>*** | ***Bih-A06-1<sup>T1</sup>, Bih-A06-1<sup>T2</sup>, Bih-A06-1<sup>T4</sup>*** | *Bih-A06-1.QN* | 10.4 | V | 88.1-95.2 |
|  | ***S-Bih-A10-1*** | ***Bih-A10-1<sup>B</sup>*** | ***Bih-A10-1<sup>T1</sup>, Bih-A10-1<sup>T2</sup>, Bih-A10-1<sup>T4</sup>*** | ***Bih-A10-1.QN*** | 21.4 | T | 43.7-46.5 |
|  | ***S-Pbr-A10-1*** | ***Pbr-A10-1<sup>B</sup>*** | ***Pbr-A10-1<sup>T1</sup>, Pbr-A10-1<sup>T2</sup>, Pbr-A10-1<sup>T4</sup>*** | ***Pbr-A10-1.QN*** | 18.8 | T | 51.7-56.7 |
|  | *S-Pbr-B06-1* | *Pbr-B06-1<sup>B</sup>* | *Pbr-B06-1<sup>T1</sup>, Pbr-B06-1<sup>T2</sup>, Pbr-B06-1<sup>T4</sup>* | *Pbr-B06-1.QN* | 4.4 | V | 176.2-184.4 |
|  | *S-Pbr-B07-1* | *Pbr-B07-1<sup>B</sup>* | *Pbr-B07-1<sup>T1</sup>, Pbr-B07-1<sup>T2</sup>, Pbr-B07-1<sup>T4</sup>* | *Pbr-B07-1.QN* | 5.7 | V | 194.1-200 |
| Days to flower | ***S-Df-A03-1*** | ***Df-A03-1<sup>B</sup>*** | ***Df-A03-1<sup>T1</sup>, Df-A03-1<sup>T2</sup>, Df-A03-1<sup>T4</sup>*** | ***Df-A03-1.QN*** | 15.3 | T | 141.3-148.2 |
|  | ***S-Df-A10-1*** | ***Df-A10-1<sup>B</sup>*** | ***Df-A10-1<sup>T1</sup>, Df-A10-1<sup>T2</sup>, Df-A10-1<sup>T4</sup>*** | ***Df-A10-1.QN*** | 40.9 | T | 53.8-56.4 |

A total of 8, 6, 5, and 3 QTL were mapped using QTL Network 2.0 for *Bf, Sf, Pf*, and *Bs*, respectively (Table S8). QTL same as the seven Stable QTL for stem strength-related traits were also detected using QTL Network 2.0 (Table S8). Significant (*p* ≤ 0.05) QTL × environment (QE) interactions were detected for the QTL *Bf* (*Bf-A10-1.QN*), *Pf* (*Pf-A10-1.QN*), and *Bs* (*Bs-A10-1.QN*) on the LG A10 (Table S8), which is consistent with the moderate heritability observed for these traits. The *Pf-A10-1.QN* and *Bf-A10-1.QN* QTL were found to be the same as the Stable QTL – *S-Pf-A10-1* and *S-Bf-A10-1*, respectively. The phenotypic variance explained by the QE interaction was low for *Pf-A10-1.QN* QTL (1.55% compared to 16.81% of the additive QTL) (Table S8). On the other hand, the phenotypic variance explained by the QE interaction for *Bf-A10-1.QN* was 6.40% compared to 13.85% of the additive QTL suggesting that this QTL might have variable R^2^ in different environments which should be taken into consideration while introgressing this QTL. The QTL detected for *Bf, Bf-A03-1.QN* showed a Type II digenic interaction with a locus on LG A01 without significant main effect. However, *Bf-A03-1.QN* QTL was not a Stable QTL for *Bf*. We did not detect any significant Type I and Type II digenic interactions among the QTL detected for *Sf, Pf*, and *Bs*.

#### Stem Diameter (*Dia*)

We identified 17 QTL for *Dia* in the TUV population using the SE trait values (Table S6), 11 of which showed positive additive effects, indicating that Tumida contributed the alleles for larger diameter, whereas six QTL were contributed by Varuna. Using BLUPs, five QTL for *Dia* were detected (Table S7). According to the adopted criteria, three QTL – one each on LGs A07, A10, and B07 were designated Stable QTL for *Dia* (Fig. 4, Table 3, and Table S10). Varuna contributed the positive allele for the major Stable QTL *S-Dia-A10-1* on LG A10 explaining 14.4% of the phenotypic variance. Tumida contributed the beneficial alleles for the Stable *Dia* QTL – *S-Dia-A07-1* and *S-Dia-B07-1* which explained 5.5% and 4.8% of the phenotypic variances, respectively.

Using QTL Network 2.0, a total of seven QTL were mapped for *Dia*, including the QTL that were the same as the three Stable QTL (Table S8). We detected significant (*p* ≤ 0.05) QE interactions for the *Dia-A10-1.QN* QTL on LG A10 which is the same as the Stable QTL *S-Dia-A10-1*. However, the phenotypic variance explained by the interaction was extremely low (Table S8) and does not have major implications for breeding efforts directed at the improvement of this trait. We did not detect any epistasis (Type I and Type II) and epistasis × environment interactions for the QTL detected for *Dia*.

#### Plant height (*Plht*)

For *Plht*, a total of 15 QTL were mapped in the TUV population using the SE trait values (Table S6). For ten of the SE QTL, Tumida contributed the alleles for higher *Plht*, which is interesting since Varuna is the better parent for *Plht*. Using BLUPs, we detected six QTL for *Plht* (Table S7). Using the adopted criteria, one QTL each on LG A06 (*S-Plht-A06-1*) and A10 (*S-Plht-A10-1*) were detected as the Stable QTL (Fig. 4, Table 3, and Table S10). The *S-Plht-A06-1* and *S-Plht-A10-1* QTL explained 6.3% and 11.3% of the phenotypic variances, respectively. The allele for shorter plant height for the QTL *S-Plht-A06-1* was contributed by Tumida, and for the QTL *S-Plht-A10-1* was contributed by Varuna.

A total of seven QTL were mapped for *Plht* using QTL Network 2.0 (Table S8); five of these QTL were also detected by Windows QTL Cartographer 2.5. One QTL each on LGs B03 (*Plht-B03-1.QN*) and B06 (*Plht-B06-1.QN*) were detected only by QTL Network 2.0. The beneficial allele in *Plht-B06-1.QN* was contributed by Tumida and therefore, it could be useful for decreasing the plant height of the Indian gene pool lines. We did not detect any QE interactions for the *Plht* QTL. Further analysis was carried out for epistasis and epistasis × environment interactions for the QTL detected for *Plht*. The QTL on LG A10 (*Plht-A10-1.QN*) exhibited Type I digenic interaction with the QTL on LG A03 (*Plht-A03-1.QN*) with additive × additive effect = −3.67 suggesting that *Plht* is decreased by the association of Varuna alleles at these loci (Table S9). The *Plht-A10-1.QN* QTL was found to be the same as the Stable QTL *S-Plht-A10-1*. Since the interaction of *Plht-A03-1.QN* and *Plht-A10-1.QN* further decreases the *Plht*, these loci need to be introgressed together.

#### Shoot branching-related traits (*Bih* and *Pbr*)

For branch initiation height (*Bih*), a total of 19 QTL were mapped in the TUV population using the SE trait values. Out of these, Tumida contributed 10 QTL for high *Bih* (Table S6). We detected four QTL for *Bih* using BLUPs (Table S7). Using the adopted criteria, a major QTL each on LG A03 (*S-Bih-A03-1*), A06 (*S-Bih-A06-1*), and A10 (*S-Bih-A10-1*) accounting for 24%, 10.4% and 21.4% of the phenotypic variances, respectively was designated as Stable QTL (Fig. 4, Table 3, and Table S10). Tumida contributed beneficial allele for low *Bih* in the *S-Bih-A06-1* QTL; therefore, this QTL would be important for improving the trait in the Indian gene pool lines of *B. juncea*.

A total of nine QTL were mapped for *Bih* using QTL Network 2.0 (Table S8). QTL same as seven of these QTL were also detected by Windows QTL Cartographer 2.5, which included the three Stable QTL for *Bih* on LGs A03, A06, and A10. *Bih-A04-1.QN* and *Bih-A09-1.QN* QTL with alleles for higher *Bih* contributed by Tumida were detected by QTL Network 2.0 only. None of the QTL detected for *Bih* showed QE and/or digenic interactions.

For the number of primary branches (*Pbr*), 16 QTL were mapped in the TUV population using SE trait values, for which both the parents, Varuna and Tumida contributed positive alleles for an increase in the number of primary branches (Table S6). We detected seven QTL for *Pbr* using BLUPs (Table S7). Stable QTL one each on LG A10 (*S-Pbr-A10-1*), B06 (*S-Pbr-B06-1*), and B07 (*S-Pbr-B07-1*) were detected for *Pbr* explaining 18.8%, 4.4%, and 5.9% of the phenotypic variances, respectively (Fig. 4, Table 3, and Table S10). Tumida contributed positive allele for a higher number of primary branches for the *S-Pbr-A10-1* QTL. Using QTL Network 2.0, a total of eight QTL were mapped for *Pbr* (Table S8). QTL same as all these QTL were also detected by Windows QTL Cartographer 2.5. The QTL controlling *Pbr* did not exhibit any QE and/or digenic (Type I and Type II) interactions.

#### Days to flowering (*Df*)

We identified 12 QTL for *Df* in the TUV population in the three trials (Table S6). Ten of these QTL were contributed by Tumida whereas two QTL were contributed by Varuna, indicating that both the parents contributed the alleles for early flowering. Using BLUPs, we detected seven QTL for *Df* (Table S7). Two QTL, one each on LG A03 (*S-Df-A03-1*) and A10 (*S-Df-A10-1*) explaining 15.3% and 40% of the phenotypic variances, respectively were designated as Stable QTL for *Df* (Fig. 4, Table 3, and Table S10); Varuna contributed the early flowering alleles in both the QTL. We detected two significant QTL each in one environment (*Df-A06-1^T2^* and *Df-A06-2^T2^*) and BLUP analysis (*Df-A06-1^B^* and *Df-A06-2B*), with Tumida contributing the early flowering alleles (Tables S6 and S7) which could explain the presence of transgressive segregants with *Df* less than Varuna in the TUV mapping population.

A total of six QTL were mapped for *Df* using QTL Network 2.0 (Table S8). QTL same as all of these QTL were also detected by Windows QTL Cartographer 2.5. We detected statistically significant (*p* ≤ 0.05) QE interactions for the *Df* QTL *Df-A10-1.QN*, which was found to be the same as the Stable QTL *S-Df-A10-1* on LG A10. The phenotypic variance explained by the *Df-A10-1.QN* QTL was 40.64% however, the phenotypic variance explained by the interaction was extremely low (1.02%) so it is inconsequential for breeding applications (Table S8). The QTL for *Df* on LG A03, *Df-A03-1.QN* (same as *S-Df-A03-1*) exhibited a Type I digenic interaction with *Df-A10-1.QN* (same as *S-Df-A10-1*) with additive × additive effect = −4.82 suggesting that *Df* is decreased by the association of Varuna alleles at both the loci (Table S9).

### Analysis of QTL clusters and conditional QTL mapping

A QTL cluster in the interval spanning 43.1-65.8 cM on LG A10 contained a Stable QTL for seven of the nine plant architectural traits mapped in the study (Fig 3 and Table S10). We detected Stable QTL with overlapping confidence intervals for *Plht (S-Plht-A10-1), Dia (S-Dia-A10-1*), and *Bih (S-Bih-A10-1*) in this cluster with Varuna contributing the allele for increasing *Dia* and decreasing *Plht* and *Bih* (Table S10). Stable QTL for stem strength-related traits, *Bf* (*S-Bf-A10-1*) and *Pf* (*S-Pf-A10-1*), with trait enhancing allele contributed by Varuna and for *Df* (*S-Df-A10-1*) and *Pbr* (*S-Pbr-A10-1*) with trait enhancing allele contributed by Tumida were also detected in the same region.

An overlap was observed in the *Pf, Bf*, and *Dia* QTL on LGs A07, A10, and B07 (Fig. 4, Table S6, and Table S7); however, it is not of major implication for stem strength improvement since the positive alleles for these traits (desirable for the improvement of stem strength) were contributed by Varuna in the colocalized QTL on LG A10 and by Tumida in the colocalized QTL on LGs A07 and B07. Similarly, the Stable QTL for *Bih* colocalized with the QTL for *Df* on LGs A03, A06, and A10 (Tables S6, S7, and S10). However, since the alleles for lower trait values were contributed by Tumida in the colocalized *Bih* and *Df* QTL on LG A06 and by Varuna in the colocalized *Bih* and *Df* QTL on LGs A03 and A10, these QTL regions can be used in breeding programs for simultaneous improvement of the two traits.

We observed an antagonistic overlap between the stable QTL for *Df* (*S-Df-A10-1*) and *Pbr* (*S-Pbr-A10-1*) on LG A10 where Varuna contributed the beneficial allele for early flowering and Tumida contributed the beneficial allele for a high number of primary branches. Similarly, the Stable QTL for *Df* on LG A03 (*S-Df-A03-1*) contributed by Tumida showed overlapping confidence intervals with a major *Pbr* QTL (*Pbr-A03-2B*) detected using BLUPs with Tumida contributing the trait enhancing alleles. These observations point toward the high correlation observed between *Pbr* and *Df* (Table 2). The regression analysis of *Pbr* on *Df* (Fig. 3) revealed several TUV lines with low *Df* and high *Pbr* suggesting that these traits are not completely pleiotropic and the high positive correlation between these traits might also be due to the linkage of some of the loci controlling them.

To identify the *Pbr* QTL without the correlated effects on *Df* and to dissect the genetic basis of control of *Df* and *Pbr* QTL on LGs A10 and A03, conditional QTL were mapped for Pbr [*Pbr*|*Df, Pbr* conditional on *Df*] that are independently expressed from *Df* (Table 4). No QTL for *Pbr* was detected on LG A10 using conditional *Pbr* values, suggesting a pleiotropic control of *Df* and *Pbr* at the *S-Pbr-A10-1* locus. However, we detected a major conditional *Pbr* QTL *Pbr-A03-1C* which was found to be the same as *Pbr-A03-2^B^* on LG A03, suggesting that this QTL is independent of the variation in *Df*; but since *Pbr-A03-2^B^* QTL was detected in only one environment and BLUP analysis, it cannot be considered a Stable QTL for *Pbr*. Additionally, four conditional *Pbr* QTL were detected on LGs A07 (*Pbr-A07-1^C^*), B03 (*Pbr-B03-1^C^* and *Pbr-B03-2^C^*), and B06 (*Pbr-B06-1^C^*) (Table 4) which could be used to improve *Pbr* in breeding programs without negative effects on *Df*.

**Table 4.**
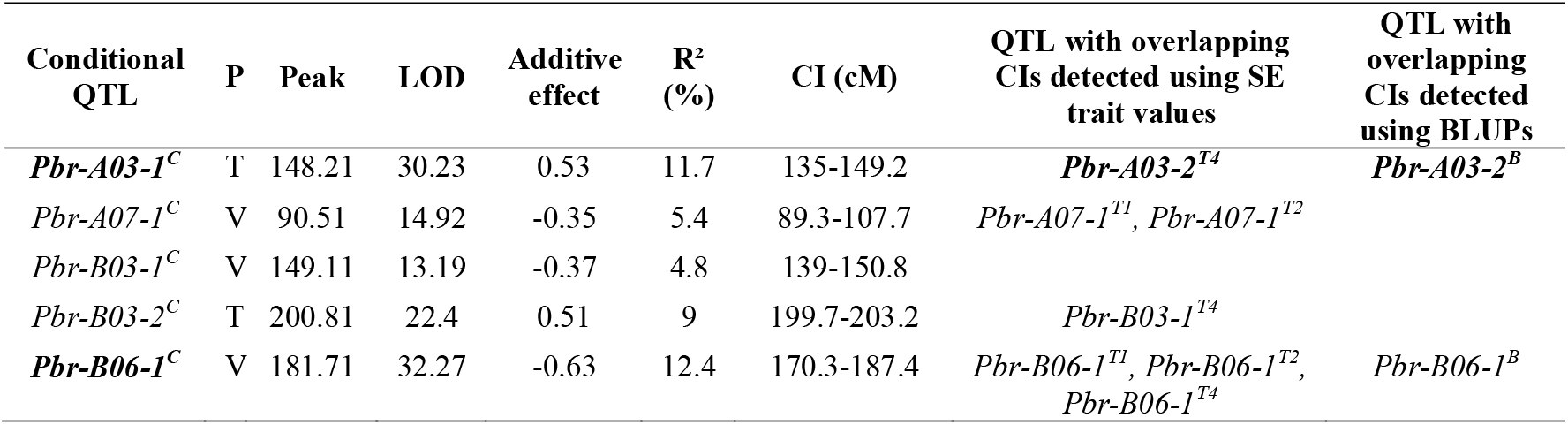
The QTL detected using conditional phenotypic values of Pbr [Pbr|Df, Pbr conditional on Df] in the TUV mapping population using Windows QTL Cartographer 2.5. Major QTL (R^2^ ≥ 10%) have been highlighted in bold. P, parent contributing the trait enhancing allele; T, Tumida; V, Varuna; R^2^, phenotypic variance explained by the QTL; CI, confidence interval; SE, single environment; T1, trial 1; T2, trial 2; T3, trial 3; T4, trial 4; BLUP, Best Linear Unbiased Prediction

### Physical intervals of the Stable QTL and identification of candidate genes

Since the genomes of both Tumida (Yang et al. 2016) and Varuna (Paritosh et al. 2021) have been sequenced, we identified the physical intervals of the Stable QTL and the number of high confidence genes in these intervals in the Varuna and Tumida genome assemblies (Tables S10 and S11). For screening the QTL intervals for the identification of candidate genes, 18 out of the 20 Stable QTL with physical intervals less than 2.5 Mb were considered. The genes contained in the Stable QTL intervals and their GO annotations are provided in Table S11. The candidate genes for each of the nine plant architectural traits were identified using the information available in heterologous systems and evidence from the functional annotations of the genes contained in the Stable QTL intervals (Table S12).

As described earlier, the transverse sections of the stems of Varuna and Tumida showed differences in interfascicular sclerenchymatous tissue (Fig. 2 and Fig. S2) therefore, the genes reported to regulate vascular cambium development and differentiation were prioritized as major candidates for the regulation of stem strength in the TUV population. Additionally, the genes with GO terms associated with biosynthesis of lignin, cellulose, and xylan were considered candidate genes for the regulation of stem strength. In the three Stable QTL for stem diameter, a total of 50 genes primarily involved in the development of stem vascular tissues, secondary cell wall biogenesis, cell growth, and auxin signaling were considered the candidate genes (Table S12).

Nine candidate genes were identified for the regulation of *Plht* in the Stable QTL *Plht-A10-1* on LG A10 (Table S12). Previous studies have highlighted the role of gibberellins, auxin, and brassinosteroids in the control of plant height (Wang and Li 2006; Wang et al. 2018; Guo et al. 2020). Therefore, the genes in the *Plht* QTL with GO terms associated with gibberellin, auxin, brassinosteroid, and strigolactone biosynthesis, transport, and signaling were also considered candidate genes for the regulation of *Plht*.

The outgrowth of axillary meristems into branches involves a complex interaction of sugars and plant hormones including auxins, cytokinins, and strigolactones; these have been well described in some reviews (Domagalska and Leyser 2011; Janssen et al. 2014; Wang et al. 2018; Barbier et al. 2019). Genes in the stable *Pbr* and *Bih* QTL with GO terms associated with shoot development, sugar transport, and signaling of plant hormones, auxins, cytokinins, and strigolactones were therefore considered as candidate genes for these traits. A total of 34 and 27 candidate genes were identified for *Bih* and *Pbr*, respectively (Table S12).

A total of 14 candidate genes for days to flowering (*Df*) were identified in the Stable QTL – *S-Df-A03-1* and *S-Df-A10-1* spanning 6.9 and 2.6 cM regions, respectively (Table S12). Out of these, 12 genes have been reported to control flowering time in literature (references listed in Table S12) and two genes have GO terms associated with the regulation of flowering time (GO:0048510, GO:2000028).

## DISCUSSION

### QTL cluster on LG A10 is critical for adaptability of the Indian gene pool lines

We detected a QTL cluster on LG A10 in the TUV population (Fig. 4) containing major loci for most of the plant architectural traits including, *Bih, Plht, Df, Bf, Pf, Dia*, and *Pbr*. This QTL cluster contains a major QTL *S-Df-A10-1* (R^2^ = 40.9%) controlling *Df* with the early flowering allele contributed by Varuna. Early flowering is the most crucial trait for the cultivation of mustard in the Indian subcontinent. QTL mapping studies in the bi-parental populations resulting from crosses between the Indian and east European lines have also revealed QTL clusters on LG A10 with the beneficial alleles for several agronomically important traits, including the flowering time, contributed by the parent belonging to the Indian gene pool (Ramchiary et al. 2007; Yadava et al. 2012).

The Stable *Df* QTL *S-Df-A10-1* in the cluster on LG A10 showed significant (*p* ≤ 0.05) digenic interaction with *S-Df-A03-1* QTL on LG A03 (Table S9). *S-Df-A03-1* and *S-Df-A10-1* contain *FT-INTERACTING PROTEIN 1* (*FTIP1*) and *CONSTANS* (*CO*) as major candidate genes, respectively. *CO* is a regulator of *FT* mRNA transcription (Samach et al. 2000) and *FTIP1* regulates *FT* protein transport (Liu et al. 2012). Since these genes are part of a common pathway for regulation of the flowering time, this could explain the digenic interaction observed between these QTL in the present study.

The alleles for low *Bih, Plht*, and *Df* and high stem strength and *Dia* for the QTL in the cluster on LG A10 are contributed by Varuna whereas, Tumida contributes the beneficial alleles for a major locus – *S-Pbr-A10-1* controlling *Pbr* in the cluster (Fig. 4 and Table 3). The confidence interval of the *S-Pbr-A10-1* locus overlaps with the *Df* locus *S-Df-A10-1* with trait enhancing alleles contributed by Tumida. QTL mapping using conditional *Pbr* values [*Pbr*|*Df*] did not detect the QTL *S-Pbr-A10-1*, suggesting a pleiotropic control of *Pbr* and *Df* at this locus and thereby, limiting the scope of its use in the improvement of *Pbr* in the Indian gene pool lines.

The regression analysis of *Pbr* with *Df* revealed several lines with low *Df* and high *Pbr* suggesting a scope of improvement of *Pbr* without increasing *Df* (Fig. 3). We identified two QTL – *Pbr-A03-1^C^* and *Pbr-B03-2^C^* in the conditional analysis (Table 4) with trait enhancing alleles from Tumida, which can be used to improve *Pbr* in the Indian gene pool lines without negative effects on *Df*.

### Engineering resistance to lodging in *B. juncea*

Stem strength is one of the key factors influencing lodging in crops (Kashiwagi and Ishimaru 2004; Ookawa et al. 2014; Wei et al. 2017; Miller et al. 2018). The four stem strength-related traits tested in the study showed moderate heritability; some significant QE interactions were observed in a few *Bf, Pf*, and *Bs* QTL (Table S8). We observed transgressive segregation for stem strength in the TUV population (Fig. S1) which was substantiated by the QTL analysis wherein both the parents have been shown to contain the QTL for increased stem strength (Tables S6 and S7). Several minor but Stable QTL for stem strength (*S-Pf-A07-1* and *S-Pf-B07-1*) were contributed by Tumida which explains the transgressive segregation observed for the trait with several TUV lines showing higher stem strength than the better parent, Varuna (Fig. S1). These QTL could be used to further improve the stem strength of the Indian gene pool lines of *B. juncea*.

Varuna, which is the better parent for stem strength, contributed the major Stable QTL for *Pf* (*S-Pf-A10-1*) and *Bf* (*S-Bf-A10-1*), both on LG A10. This is consistent with the observation of a reduced number of interfascicular sclerenchymatous cell layers in the transverse stem sections of Tumida (Fig. 2). A correlation of reduced stem vascular tissues with stem lodging has also been previously reported in the *MYB43* RNAi lines of *B. napus* (Jiang et al. 2020). The stem strength QTL with beneficial alleles from Varuna could be used to improve the good combiners of hybrids in the east European gene pool with low stem strength, e.g., *B. juncea* lines EH-2 and S7 (Table S13). The Indian gene pool lines of *B. juncea* have a narrow genetic diversity (Srivastava et al. 2001; Burton et al. 2004), however, we observed high variability in the stem strength-related traits within the Indian gene pool lines (Table S13). The stem strength QTL – *S-Pf-A10-1* and *S-Bf-A10-1* on LG A10 with beneficial alleles contributed by Varuna can also be used to improve the trait in the Indian gene pool lines having low stem strength.

We observed a high correlation of stem strength-related traits (*Pf* and *Bf*) with stem diameter (*Dia*) (Table 2). A high correlation between absolute stem strength and stem diameter has also been reported in *B. napus* (Miller et al. 2018). An overlap was observed in the confidence intervals of the Stable QTL identified for these traits on LGs A07, A10, and B07 (Fig. 4, Table S6, and Table S7). Based on these results, a pleiotropic control of stem strength and stem diameter could be hypothesized at these loci. Since the beneficial alleles for both *Pf* and *Dia* in the QTL with overlapping confidence intervals on LGs A07 and B07 are contributed by Tumida, these loci could be used to simultaneously improve stem strength and diameter in the Indian gene pool lines. A Stable locus for *Dia* (*S-Dia-A10-1*) was identified in the QTL cluster on LG A10 with the beneficial allele from Varuna. The locus contains *CSLD2* and *PIN5* genes that have been previously implicated in the control of stem diameter in *Arabidopsis* and *Populus* (Yin et al. 2011; Zheng et al. 2021).

Most of the mustard cultivars grown in India have high *Bih* with branches initiating up to 1m from the ground. These lines are more prone to top lodging under stormy conditions at maturity due to top-heavy branches resulting in substantial yield losses (Chakrabarty et al. 1994). Therefore, the reduction of *Bih* is a major objective for mustard improvement programs in India. *Bih* showed a significant positive correlation with *Df* and *Plht* in the TUV population (Table 2). This is consistent with a previous study in *B. napus* which showed a significant positive correlation and co-localization of the QTL for the two traits (Shen et al. 2018). However, Vijayakumar et al. (1996) reported a negative correlation of *Bih* with plant height and flowering time in *B. juncea*. We observed a high positive correlation (Table 2) and an overlap between the QTL for *Bih* and *Df* (Tables S6 and S7). The overlap between the Stable *Bih* and *Df* QTL on LGs A10 and A03 (Fig. 4) is not of major consequence for breeding since the alleles reducing both *Bih* and *Df* are contributed by Varuna in the overlapping QTL. The overlapping Stable QTL for *Bih* (*S-Bih-A03-1*) and *Df* (*S-Df-A03-1*) on LG A03 contained the *TFL1* gene (Table S12) that has been previously implicated to pleiotropically control branch number, *Df* and branch initiation height in *B. napus* (Sriboon et al. 2020). Tumida contributed the allele for reduction of *Bih* in *S-Bih-A06-1*, therefore, this locus can be introgressed in the Indian gene pool lines to reduce the branch initiation height.

The use of dwarfing genes has been shown to improve the resistance to plant lodging in *B. napus* (Muangprom et al. 2006; Liu et al. 2010; Yang et al. 2021). Therefore, the introgression of *the S-Plht-A06-1* locus from Tumida could further improve the lodging resistance of the Indian gene pool lines via reduction of *Plht*. Though Tumida shows a low trait value for *Plht* as compared to Varuna, it contributed a major QTL *S-Plht-A10-1* for the enhancement of the trait. Significantly, no overlap was observed in the QTL identified for stem strength and *Plht*, which provides an opportunity to simultaneously introgress beneficial loci for these traits to tackle the problem of plant lodging.

### Conclusions

We used a high-density linkage map to identify 20 Stable QTL for nine plant architectural traits related to stem strength, stem diameter, plant height, shoot branching, and days to flowering in the TUV F_1_DH population. The Chinese stem type mustard, Tumida contributed positive alleles for all the nine architectural traits and could be a novel source for beneficial alleles for further improvement of these traits in the Indian gene pool lines. The QTL cluster on A10 contains a major locus for *Df* and is critical for the early flowering of the Indian gene pool lines therefore, no introgressions can be made in the Indian gene pool lines in this region. The study has identified conditional QTL for *Pbr* with trait enhancing alleles contributed by Tumida, which could be used to improve the trait without negative effects on *Df*. The QTL for *Pf, Bf, Sf, Dia, Bih*, and *Plht* identified in the study could be used for the genetic improvement of lodging resistance in *B. juncea* by the enhancement of stem strength in addition to an increase in stem diameter and a reduction in branch initiation height and total plant height. The Stable QTL for plant architectural traits could be introgressed into pure lines or combiners of hybrids that lack the superior alleles for the mapped architectural traits.

Altogether, the findings reported in the study have revealed the genetic control of some of the key plant architectural traits and would find application in breeding for improved ideotypes in *B. juncea*.

## Statements & Declarations

### Author Contributions

PS, SM, and VG carried out the phenotyping. SM, PS, and SKY carried out the data analysis and mapping work. SM performed the stem anatomy and candidate gene analyses. DP and AKP conceived and supervised the overall study. SM and DP drafted the manuscript. All the authors reviewed and approved the final draft of the manuscript.

## Acknowledgements

We thank Satish Giri for his help with the field experiments. Microscopy was carried out at the Central Instrumentation Facility, University of Delhi South Campus. The research was supported by the Department of Biotechnology (DBT), Government of India, under the project ‘DBT-UDSC partnership Centre on Genetic Manipulation of Brassicas’, grant number – BT/01/NDDB/UDSC/2016. SM was supported by a research fellowship (Ref. no. 19/06/2016(i)EU-V) from the University Grants Commission (UGC).

## Data Availability

The datasets generated during and/or analyzed during the current study are available from the corresponding author on request.

## Ethical standards

The experiments were performed in compliance with the laws of India.

## Supplementary Material

### Supplementary figures

**Fig. S1** The density histograms of TUV population for bend force (*Bf*), stab force (*Sf*), press force (*Pf*), breaking strength (*Bs*), stem diameter (*Dia*), plant height (*Plht*), branch initiation height (*Bih*), number of primary branches (*Pbr*), and days to flower (*Df*). Blue, red, and green arrows indicate Tumida, Varuna, and TUV-F_1_, respectively. The Y-axis represents the density (the ratio of frequency to group distance) for each trait

**Fig. S2** Anatomical and phenotypic differences in Tumida, Varuna and some strong (s) and weak (w) TUV population lines sampled at mature green stage. (a-f) Cross sections of midpoint of last internode showing differences in layers of interfascicular sclerenchyma tissue (depicted by yellow lines). Scale bars represent 50 μm. g. Phenotypic differences (BLUPs) in stem diameter (*Dia*) and stem strength measures (*Bf*, bend force; *Bs*, breaking strength; *Sf*, stab force; *Pf*, press force)

### Supplementary tables

**Table S1** Details of the trials conducted for phenotyping plant architectural traits in the TUV F_1_DH population

**Table S2** Distribution and density of GBS markers on the TUV linkage map

**Table S3** Positions of the GBS markers on the TUV linkage map

**Table S4** Trait statistics and broad sense heritability (H_B_) of plant architectural traits in TUV F_1_DH population in different phenotypic trials (T1, T2, T3, and T4). *Bf, Sf, Pf, Bs, Dia, Plht, Bih, Pbr*, and *Df* represent bend force, stab force, press force, breaking strength, stem diameter, plant height, branch initiation height, number of primary branches, and days to flower, respectively

**Table S5** Trait values (BLUPs) for plant architectural traits in TUV F_1_DH mapping population (169 lines). *Bf, Sf, Pf, Bs, Dia, Plht, Bih, Pbr*, and *Df* represent bend force, stab force, press force, breaking strength, stem diameter, plant height, branch initiation height, number of primary branches, and days to flower, respectively

**Table S6** Single environment (SE) QTL detected for all traits in TUV F_1_DH mapping population (169 lines) using Windows QTL Cartographer 2.5. Major QTL (R2 ≥ 10%) have been highlighted in bold. R^2^, phenotypic variance explained by the QTL; T1, trial 1; T2, trial 2; T3, trial 3; T4, trial 4; T, Tumida; V, Varuna

**Table S7** QTL detected using BLUPs for nine traits in the TUV F_1_DH mapping population (169 lines) using Windows QTL Cartographer 2.5. Major QTL (R2 ≥ 10%) have been highlighted in bold. R^2^, phenotypic variance explained by the QTL; T1, trial 1; T2, trial 2; T3, trial 3; T4, trial 4; T, Tumida; V, Varuna

**Table S8** The main effect QTL detected by QTL Network 2.0 in the TUV F_1_DH mapping population. The main effect QTL also independently detected by Windows QTL Cartographer 2.5 are highlighted in bold. The QTL showing significant (p ≤ 0.05) QTL × environment interactions are highlighted in blue. A, additive effect; AE, additive effect due to QTL-environment interaction; h^2^(a), phenotypic variance explained by the QTL; h^2^AE, phenotypic variance explained by the QTL × environment interaction; T1, trial 1; T2, trial 2; T3, trial 3; T4, trial 4; T, Tumida; V, Varuna; *Bih*, branch initiation height; *Df*, days to flower; *Pbr*, number of primary branches; *Plht*, plant height; *Dia*, stem diameter; *Bf*, bend force; *Sf*, stab force; *Pf*, press force; *Bs*, breaking strength

**Table S9** Epistatic interactions between QTL detected by QTL Network 2.0 for plant architectural traits in TUV F_1_DH mapping population (169 lines). The main effect QTL are highlighted in bold. e, environment; T1, trial 1; T2, trial 2; T3, trial 3; T4, trial 4; T, Tumida; V, Varuna; *Bih*, branch initiation height; *Df*, days to flower; *Pbr*, number of primary branches; *Plht*, plant height; *Dia*, stem diameter; *Bf*, bend force; *Sf*, stab force; *Pf*, press force; *Bs*, breaking strength

**Table S10** Stable QTL for plant architectural traits in the TUV population. The major QTL (R^2^ ≥ 10%) are highlighted in bold. T, Tumida; V, Varuna; SE, single environment; T1, trial 1; T2, trial 2; T3, trial 3; T4, trial 4

**Table S11** List of genes harbored in the Stable QTL regions (< 2.5Mb) for plant architectural traits in the TUV population according to the Tumida and Varuna genome assemblies

**Table S12** Candidate genes in the Stable QTL intervals (< 2.5Mb) for plant architectural traits in the TUV population

**Table S13** Mean (n = 10) trait values of the stem strength measures of some *B. juncea* cultivars calculated using the phenotypic data obtained in the field trial conducted in 2019-20 (T2)

