## Supplementary Figures for "Genetic mapping of some key plant architecture traits in *Brassica juncea* using a cross between two distinct lines – vegetable type Tumida and oleiferous Varuna"

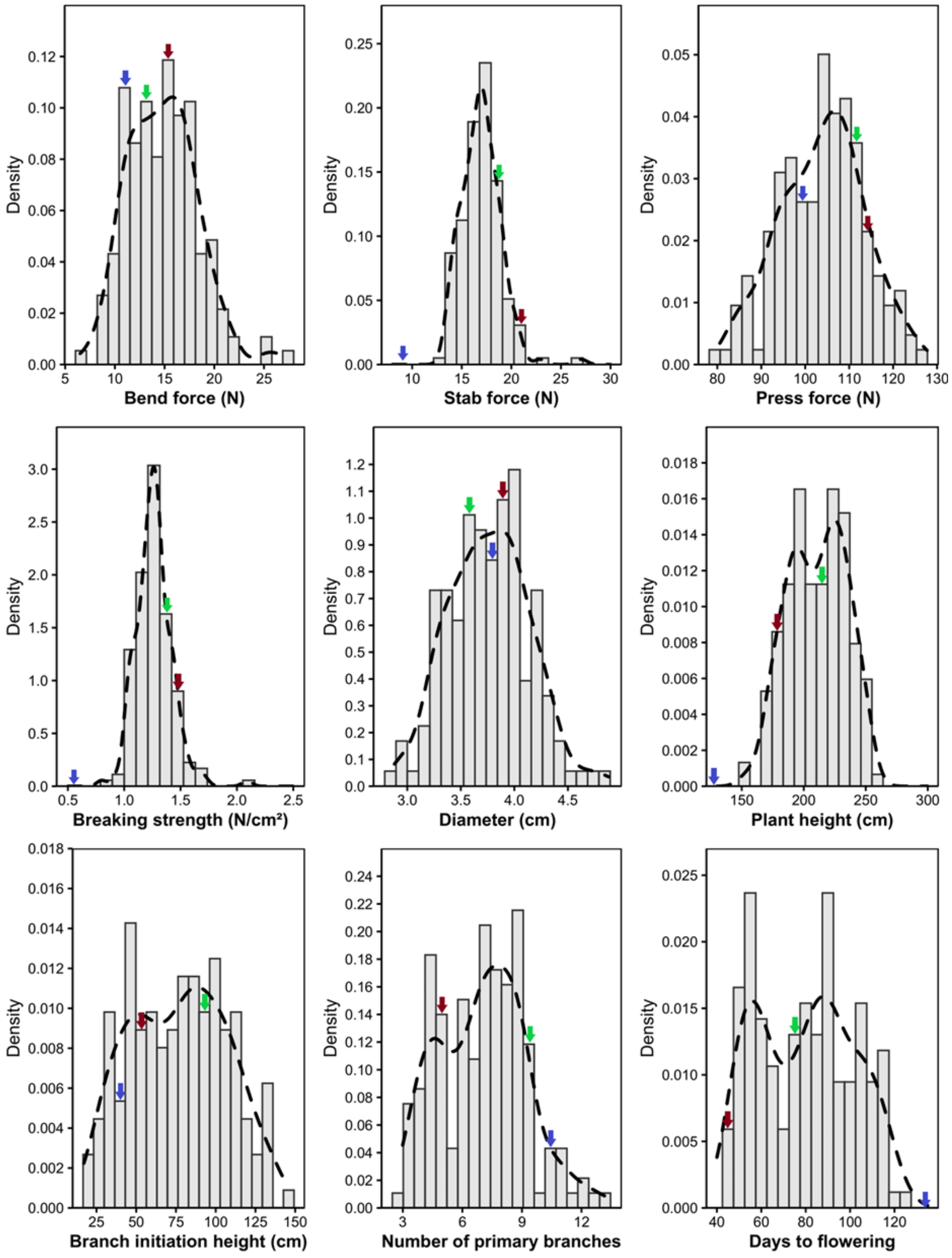


**Fig. S1** The density histograms of TUV population for bend force (*Bf*), stab force (*Sf*), press force (*Pf*), breaking strength (*Bs*), stem diameter (*Dia*), plant height (*Plht*), branch initiation height (*Bih*), number of primary branches (*Pbr*), and days to flower (*Df*). Blue, red, and green arrows indicate Tumida, Varuna, and TUV-$F_1$, respectively. The Y-axis represents the density (the ratio of frequency to group distance) for each trait

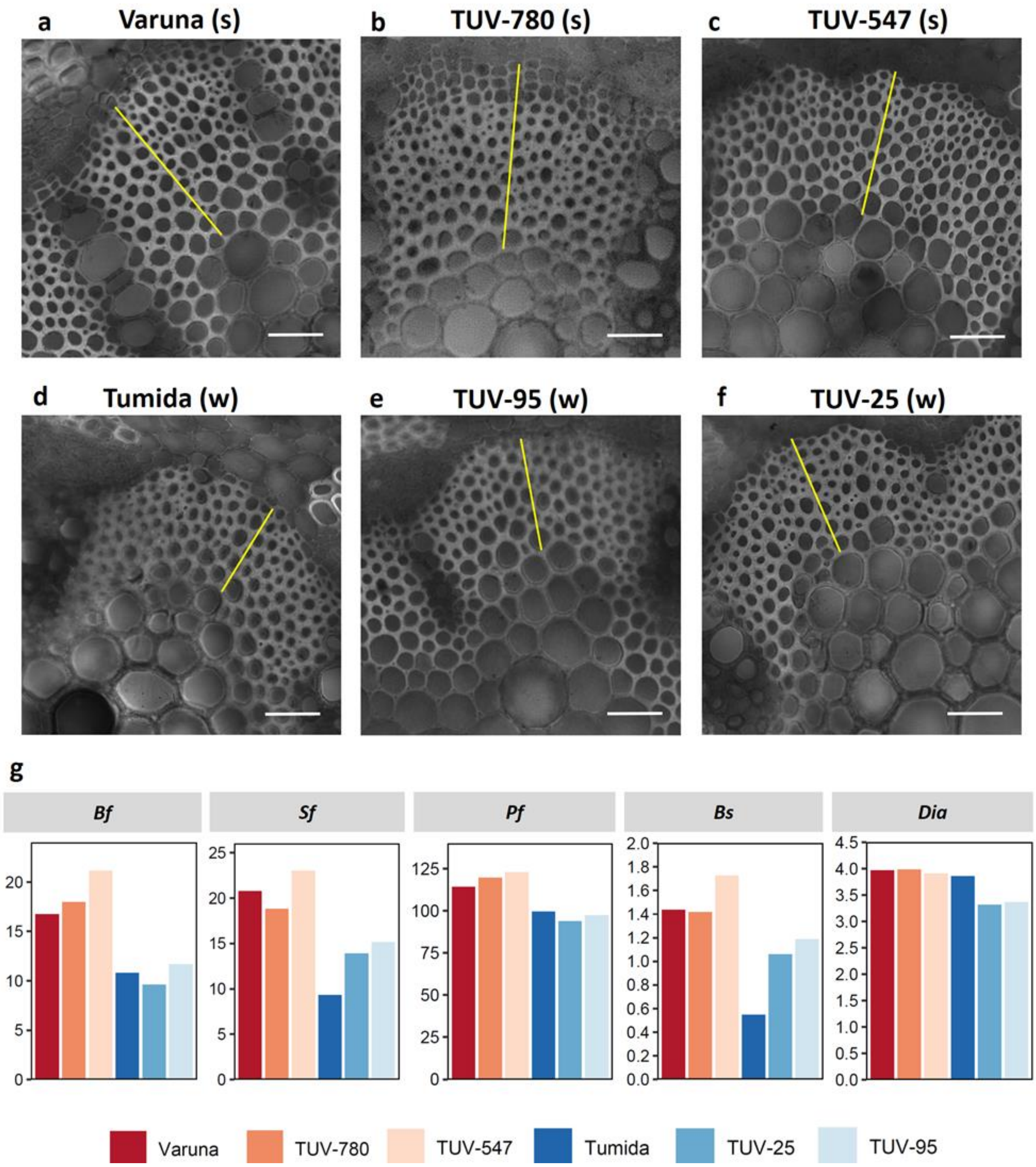


**Fig. S2** Anatomical and phenotypic differences in Tumida, Varuna and some strong (s) and weak (w) TUV population lines sampled at mature green stage. (a-f) Cross sections of mid-point of last internode showing differences in layers of interfascicular sclerenchyma tissue (depicted by yellow lines). Scale bars represent 50 μm. g. Phenotypic differences (BLUPs) in stem diameter (*Dia*) and stem strength measures (*Bf*, bend force; *Bs*, breaking strength; *Sf*, stab force; *Pf*, press force)
